# Testosterone Administration Modulates Competitive Choice via Ventral Striatum and TPJ

**DOI:** 10.64898/2026.05.05.722878

**Authors:** Jan B. Engelmann, Veerle van Son, Karin Roelofs, Alan Sanfey, Ale Smidts, Pranjal Mehta

## Abstract

How does testosterone influence decisions and choice-related neural computations in competitive environments? To address this question, we administered testosterone or placebo to female participants (n = 54) in a double-blind, randomized design. Following drug treatment, participants competed in a dot estimation task that manipulated opponent status (lower, equal, or higher) and outcome feedback (win or loss), after which they decided whether to compete against the same opponent again. All participants adjusted their behavior based on opponent status and outcome feedback. Participants who received testosterone, however, showed significantly greater sensitivity to outcome feedback: they were more willing to compete after winning and less willing after losing, and made those decisions faster - suggesting that testosterone increases the weighting of immediate, salient outcome information in competitive decision-making. At the neural level, a network comprising ventral striatum, vmPFC, bilateral TPJ and ACC processed outcome-related signals during the feedback period. Critically, neural prediction analyses at the trial-level revealed that activity in left ventral striatum and TPJ predicted subsequent decisions to compete, but only in participants who received testosterone. The direction of these effects mirrored the behavioral results: striatal activity amplified the tendency to re-compete after winning, whereas TPJ activity predicted renewed competition after losing. Together, these findings demonstrate that testosterone biases competitive decisions by amplifying the influence of outcome-related activity in reward and social cognition circuits.

## Introduction

Testosterone is a steroid hormone that modulates social behavior. While initial reports of increased aggressive behavior have shaped much of folk theory about the effects of testosterone, more recent evidence indicates that the effects of testosterone are highly nuanced and in fact depend on the social context. One influential theory states that testosterone is associated with the pursuit of dominance, and therefore influences status seeking behavior in competitive environments (Eisenegger, Haushofer, and Fehr 2011; Mazur and Booth 1998). Indeed, non-human and human experiments have provided evidence consistent with this notion: both endogenous and exogenous testosterone are positively associated with competitive behavior concerning social status and dominance (Beehner et al. 2006; Cashdan 1995; Mazur and Booth 1998; Sapolsky 1991).

While a growing subset of behavioral studies have addressed the causal effect of exogenous testosterone administration on status seeking behavior (Dreher et al. 2016; Losecaat Vermeer et al. 2020; Mehta et al. 2015; Nave et al. 2018; but see Polo et al. 2024), the effect of testosterone on the neural correlates of such behavior has been studied very little to date. The current study aimed to address this gap in the literature by identifying the causal impact of testosterone on the neural correlates of competitive behavior in a social competition task that specifically manipulated context (Mehta et al. 2015). In particular, two aspects of the social context were manipulated: (1) whether participants competed against an opponent of higher, equal or lower status (based on ability) and (2) whether a prior dominance contest resulted in victory or defeat (Mehta et al. 2015; Mehta, Jones, and Josephs 2008). These contexts underlie status seeking behavior in competitive environments, and manipulating them directly in the current study allowed us to investigate testosterone’s context-dependent influences on competitive decisions and their neural correlates.

Previous research on the neural effects of testosterone administration has demonstrated consistently increased activity in ventral striatum and amygdala in social-emotional tasks (Hermans et al. 2010; Radke et al. 2015). A closer look at the relatively small number of fMRI studies included in a recent meta-analysis (Heany et al. 2016), however, reveals that the effects of testosterone administration on neural activation patterns are highly task-dependent. Specifically, previous research has demonstrated that testosterone increased activation in ventral striatum during reward anticipation in the Monetary Incentive Delay (MID) task (Hermans et al. 2010), consistent with the role of the ventral striatum in processing reward-related information (Knutson and Greer 2008; Pessoa and Engelmann 2010). Enhanced amygdala activation due to testosterone administration, on the other hand, is usually observed in tasks requiring the processing of emotional information from faces, such as their trustworthiness (Bos et al. 2012), or emotional expressions of fear, anger and happiness (Bos et al. 2013; Radke et al. 2015). Jointly these results suggest that testosterone modulates neural circuitry that supports ongoing tasks in important ways, particularly for reward processing, emotion identification and emotional action control.

Notably little is known about the modulatory influences of testosterone on the neural correlates of decision making in social contexts (but see (Bos et al. 2012; Ou et al. 2021). In our competitive choice task, two neural circuits are of particular relevance: 1. social cognition regions and 2. valuation, or reward-related regions. Social cognition regions, such as the temporoparietal junction (TPJ) and dorsomedial prefrontal cortex (dmPFC), have consistently been implicated in simulating and understanding other peoples’ mental states, a computation commonly referred to as mentalizing or theory of mind (Carter and Huettel 2013; Chang et al. 2023; Van Overwalle 2009). Understanding competitors’ relative status in relation to one’s own and translating this information into a competitive decision is fundamentally a social cognitive computation, which we hypothesized to be processed by regions in the social cognition network (Chang et al. 2023; Muscatell et al. 2012; Zink et al. 2008). Moreover, our participants experienced victory or defeat against an opponent before they made decisions to compete. Such outcome-related information is commonly processed by the brain’s valuation network, consisting of ventral striatum and ventromedial prefrontal cortex (Bartra, McGuire, and Kable 2013; Pessoa and Engelmann 2010). This same network also processes social value (Ruff and Fehr 2014), and therefore fundamentally supports evaluations of the social outcomes presented on every trial in our experiment. Given the task-dependent effects of testosterone on neural activation patterns demonstrated in prior studies (Heany et al. 2016), together with the strong involvement of valuation regions in processing social outcomes and social cognition regions in computing relative social standing, we expected testosterone to modulate outcome-related information in VS and VMPFC, as well as social cognition regions in TPJ, as these computations jointly support competitive decision-making in the current task.

## Methods

### Participants

A total of 54 healthy female subjects participated in the experiment (Age = 21.6, SD 2.4), all of whom gave their informed consent to experimental procedures that were approved by the Radboud University ethics committee. Participants reported no history of psychiatric, neurological, or endocrine disease and were not currently using corticosteroids. Only women were included in the present experiment because the pharmacokinetics of our testosterone administration procedure have only been established in women and not yet in men at the time of the study (Tuiten et al. 2000), Moreover, this decision is in keeping with the bulk of prior testosterone administration studies that have adopted this same dosage and sublingual administration technique (Bos et al. 2012; Eisenegger et al. 2010; Hutschemaekers et al. 2021, 2023; Radke et al. 2015).

Participants who displayed little to no choice variability (less than 10% of choices in one decision category (compete/no compete) across at least 2 runs, N = 8) were excluded from all analyses, because (1) imaging models including parametric modulators cannot be estimated with such little choice variance, (2) such a low number of events in one category leads to reduced detection power (Desmond and Glover 2002), particularly when using thresholding methods based on spatial extent (Huettel and McCarthy 2001) and (3) such perseverating choice behavior might be indicative of having misunderstood the task, or using an idiosyncratic choice strategy. The final sample that was included in all subsequent analyses therefore included a total of 46 subjects (Placebo group N = 22; Testosterone group N = 24).

### Procedure

#### Testosterone Administration and Hormone Assays

Testosterone administration and hormone assays followed established procedures outlined in detail in Mehta et al. (2015). Briefly, in a double-blind between-subjects design, participants self-administered a single sublingual dose of either testosterone (0.5 mg testosterone, 0.5 mg hydroxypropyl-cyclodextrin, 0.005 mL 96% ethanol, and distilled water) or placebo. This dosage has been shown to increase serum testosterone levels tenfold within 15 minutes, with effects peaking 4-6 hours post-administration (Bos et al. 2012; Eisenegger et al. 2010; Tuiten et al. 2000). We therefore started behavioral testing approximately 4.5 hours after administration. Saliva samples, analyzed using a double-antibody luminescence immunoassay kit (at Clemens Kirschbaum’s laboratory in Germany), showed no significant baseline testosterone differences between groups (testosterone: M = 22.11 pg/mL, SD = 11.74; placebo: M = 22.52 pg/mL, SD = 15.00, t(49) = −0.11, p = 0.92), with average inter-assay and intra-assay coefficients of variation at 6.48% and 8.56%, respectively.

#### Experimental Task

The experimental task is illustrated in Figure 1A and outlined in detail in Mehta et al, (2015). Briefly, participants performed a competition task with a repeated one-shot design while lying in the MRI scanner. They were told that they would compete against 90 other women who had completed the task earlier and acquired a status that was reflective of their prior performance in the dot estimation task. Status was signified by stars, with one star signifying low performance, two stars intermediate (the same as the subject in the scanner), and three stars high performance. In fact, however, those opponents were simulated and winning or losing was predetermined, with 50% of trials resulting in win/loss for the pairings with players from each status category. Participants were assigned to the two-star group, and were told that their status would be updated at the end of the task. Status of the opponent (one, two, or three stars) and the participant’s outcome feedback for each trial (win or lose) were experimentally manipulated in a 2 (win/loss) x 3 (lower (*), equal (**), higher (***) status opponent) repeated measures design. Importantly, after competing on a dot estimation problem, and finding out whether they won or lost, participants were asked whether they wanted to compete again against this particular opponent or preferred a dot estimation trial without an opponent. A trial without an opponent would not count towards their ranking. Participants were told that the re-challenge trial or the trial without an opponent would take place after participants came out of the scanner, but never actually occurred.

**Figure 1.**
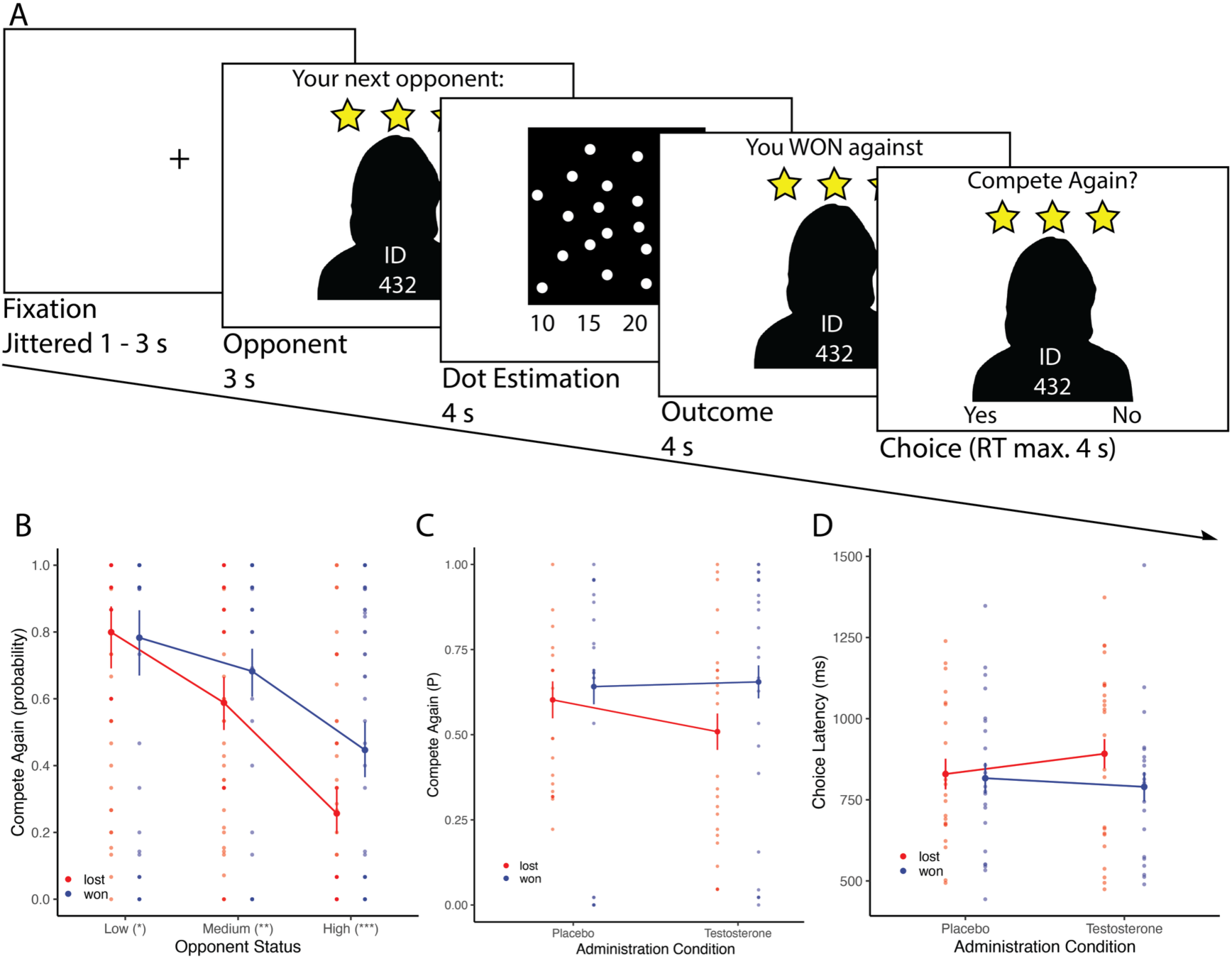
Task and Behavioral Results. (A) Task schematic and timing. After a fixation cross was presented for a variable period between 1 and 3 seconds, participants were informed of the opponent status for the current trial (3 seconds). They provided their estimate on the dot estimation problem within a time window of 4 seconds and were informed whether they won or lost against their opponent. Finally, they decided whether to compete again or not at the end of the trial within a period of 4 seconds. B-D show results from trial-by-trial mixed regressions. Outcome and Opponent Status jointly influenced subsequent decisions to compete (B). Moreover, (C) shows that testosterone administration moderated the influence of Outcome on decisions to compete, indicating that participants that received testosterone were more sensitive to winning and losing than Placebo subjects, and (D) made those decisions relatively faster after winning. Error bars reflect model-based estimates of the standard error, dots reflect subject-wise means per condition based on the raw data.

Trials started with a fixation cross for 1000, 2000 or 3000 ms, followed by information about the opponent (ID number and status) for 3000 ms. A dot estimation problem was then presented for 4000 ms, and participants were asked to enter their estimate of the number of dots on screen by pressing one of four buttons. The outcome screen, presented for 4000 ms, provided participants with ‘won’ or ‘lost’ feedback, and reminded them of their opponent’s status for this trial. Participants had a response window of 4000 ms for the decision to compete again. The task was divided into 3 runs with 30 trials each. E-prime was used to present stimuli and record responses and reaction times. If no response to the dot estimation problem was detected, loss feedback was shown (0.006% of all trials) and this trial was excluded from further analyses.

#### Behavioral analysis approach

A separate analysis of these behavioral data focusing on individual differences was reported previously (Mehta et al. 2015). In contrast, the present analyses differ in three key ways: (1) they emphasize trial-by-trial behavior rather than individual differences, (2) they use mixed-effects models that leverage trial-level observations to estimate predictors of choice while accounting for between-participant variability, and (3) they apply different exclusion criteria, optimized here to ensure reliable regressors for the fMRI analyses (see Participants section). Decisions to compete (not compete = 0, compete = 1) and reaction times were analyzed on a trial-by-trial basis using a generalized linear mixed-effects model that was implemented via the *mixed* function of the R package afex (Singmann and Kellen 2019), which calls the *glmer* and *lmer* functions of the lme4 package (Bates et al. 2015). A maximal random-effects structure was initially specified, including a participant-specific random intercept, random slopes for all within-subject predictors, and correlations among all random effects (Barr 2013). To address convergence issues, we simplified the structure stepwise: first by removing correlations among random effects, and subsequently by removing or simplifying random slopes as needed (Matuschek et al. 2017). Specifically, we dropped the Outcome random slope and modeled Status as a continuous predictor rather than a factor to improve stability. For all analyses (choices, choice latencies and neural prediction analyses), we report results from the most complex model that converged, typically including random intercepts for subjects and random slopes for Status. The final model structures are detailed below and in Table S1. *p*-values were obtained from likelihood ratio tests (LRT). All continuous predictors were standardized and categorical predictors sum-to-zero coded. Follow-up tests were conducted using the *emmeans* package (Lenth 2016), which estimated marginal means and simple slopes from the fitted model; log-odds estimates were then exponentiated to obtain odds ratios with corresponding 95% confidence intervals (reported in brackets).

#### fMRI Acquisition

Images were acquired on a 1.5 T Siemens Avanto MRI scanner (Siemens Medical Systems, Erlangen, Germany) equipped with an eight-channel head coil. High-resolution anatomical images were obtained using a T1-weighted MP-RAGE sequence (TR = 2250 ms, TE = 2.95 ms, TI = 850 ms, flip angle = 15°, 176 sagittal slices, voxel size = 1.0 × 1.0 × 1.0 mm³, FoV = 256 mm). Functional images were acquired using a multi-echo EPI sequence with GRAPPA parallel imaging (Poser et al. 2006) (TR = 2140 ms, TEs = 9.4/21/33/44/56 ms, flip angle = 90°, 34 axial slices, ascending acquisition, slice thickness = 3.0 mm, distance factor = 17%, in-plane voxel size = 3.3 × 3.3 mm², FoV = 212 mm, GRAPPA factor = 3).

#### fMRI Analyses

Preprocessing and statistical analyses were performed using SPM12 (Department of Imaging Neuroscience, UCL, London, UK). The first two volumes of the second, and third functional runs were discarded for T1 equilibration effects. Based on the 30 prescans of the first run, optimal weighting parameters for each of the five echo times were calculated per participant, which were then used to combine the five echoes into a single volume (Poser et al. 2006). To correct for head motion, all functional volumes were realigned to the first volume using septic b-spline interpolation, followed by slice timing correction to the middle slice. To improve coregistration, bias-corrected anatomical and mean EPI images were created and subsequently coregistered using segmentation. Images were normalized to the Montreal Neurological Institute T1 template using the parameters (forward deformation fields) derived from the nonlinear normalization of individual gray matter tissue probability maps. Finally, functional data underwent spatial smoothing using an isotropic 8-mm FWHM Gaussian kernel. Although slice-timing correction was applied to account for ascending acquisition, residual uncertainties in the alignment between behavioral event logs and fMRI time series are well documented (Parker and Razlighi 2019; Viswanathan et al. 2020). To address this, we estimated three models with regressors shifted by 0, +1, and +3 s. Our main findings were robust across these models (see Figure S1). We report results from the intermediate model that implemented a +1 s shift.

The first-level model included regressors modeling the four different periods encountered by subjects throughout the task (see **Figure 1A**), including the (1) opponent information period, (2) dot estimation task, (3) outcome information period, and (4) decision period. To capture differential activation depending on subjects’ decisions, the last period was binned by the type of decision made by subjects (compete vs. no compete). Decisions during the dot estimation task and compete choices were modeled as boxcars covering the period from trial onset until the button press indicating choice (variable epoch model; (Grinband et al. 2008; Yarkoni et al. 2009) to best capture the exact decision period, minimize distortions due to time on task effects, and implicitly model choice latencies. All other periods were modeled with delta functions of length 0. Opponent status was modeled as a parametric modulator throughout all the stages of the task. Finally, six movement parameters and the average global signal (extracted from all voxels within the SPM mask) were included as regressors of no interest and to account for residual head-movement related and other sources of noise (Power, Schlaggar, and Petersen 2015).

We focus our fMRI analyses on outcome- (reflecting whether participants won/lost the competition) and choice-related activity (reflecting whether subject chose to compete again or not), as the neural signatures during these periods most closely relate to our behavioral analysis of the impact of outcome on decisions to compete. Contrasts of interest were performed to test activations patterns during the outcome and decision periods. Specifically, during the outcome period we tested both the contrast winning vs losing and the outcome event independent of having won or lost, while during the decision period we tested the contrast compete vs. no compete and the decision event independent of the decision made. Moreover, during these periods, we inspected to what degree activation patterns depended on the opponent status by performing all contrasts mentioned above on the parametric modulator reflecting status. For all contrasts we inspected the data for average activation patterns that were consistent across all subjects, as well as the differential effects of testosterone administration. All results are corrected at the cluster level at an FWE-corrected threshold of p<0.05 (initial cluster forming height threshold of p<0.001).

#### Neural Prediction Analyses

To identify whether signal in outcome relevant regions predicts overt choices to compete during the subsequent (4 seconds later) decision period, we extracted signal from valuation and social cognition regions of interest that showed significant activation during the outcome period (**Table S2**). The goal of these analyses was to identify brain-behavior relationships by regressing this trial-by-trial regional signal during the outcome period against the subsequent decisions made by subjects. This approach allowed us to test (1) whether regional signal predicts subsequent decisions after winning compared to after losing, (2) whether regional signal differentially relates to subsequent choices when interacting with opponents of different status levels, and (3) whether signal is differentially related to decision-making as a function of testosterone administration.

Activation time courses were extracted from 6mm spherical volumes of interest (VOIs) centered on the activation foci listed in **Table S2**. For each participant, the first eigenvariate of the adjusted BOLD signal was extracted from each VOI, using the SPM12 VOI toolbox. This procedure yields a representative time series of regional activity that is adjusted for effects of no interest, rather than contrast estimates. To reduce the influence of extreme signal fluctuations, values exceeding the mean by more than 3 standard deviations were replaced with the maximum value within the range of mean ± 3 SD (“despiking”). Data were spatially averaged over the sphere, and were temporally downsampled to one value for each trial, by averaging over 3 images within the period of 6.4s (3TRs) – 10.7s (5TRs) after trial onset. This ensured that the data summarized the delayed and dispersed hemodynamic response reflecting outcome- and choice-relevant computations. Spatial and temporal averaging and correction for session and movement confounds were implemented using the VOI toolbox in SPM12. We implement trial-by-trial analyses of BOLD signal extracted from VOIs via generalized linear mixed-effects models (GLME) using the *mixed* function of the R package afex.

As with our behavioral models, we initially specified a maximal random-effects structure for the neural models and applied the same procedure to address convergence issues. These models are inherently more complex than the behavioral models due to the inclusion of the neural signal, and mixed *logistic* models in particular are prone to convergence problems when fit with maximal random-effects structures, as binary outcomes provide less information per observation and the logit link results in a more complex likelihood surface (Bates et al. 2015; Bolker et al. 2009; Matuschek et al. 2017). We therefore simplified the random-effects structure stepwise: first by removing correlations among random effects, and then by removing or simplifying random slopes as needed (Matuschek et al. 2017). Specifically, we dropped the Outcome random slope and modeled Status as a continuous predictor rather than a factor across all models; in some cases, we also removed correlations among the remaining random slopes. For each analysis, we report results from the most complex model that converged and specify the exact random-effects structure in Tables S4 and S5.

#### Behavioral Results

Decision results are shown in Table S1 (left column) and indicate main effects of opponent Status (χ²(2) = 31.46, p < 0.001) and trial Outcome (won vs. lost; χ²(1) = 27.81, p < 0.001). In contrast, we observed no main effect of Testosterone Administration (χ²(1) = 0.28, p = 0.598). Moreover, a significant Outcome × Status interaction (χ²(2) = 26.99, p < 0.001) showed that participants responded differentially to won–lost outcomes depending on their opponent’s status. Specifically, as shown in **Figure 1B**, in their subsequent decisions to compete they did not differentiate between won and lost trials for low-status opponents (OR = 1.11 [0.83 1.47], p = 0.46), but did so for intermediate- (OR = 0.67 [0.57 0.79], p < 0.001) and high-status opponents (OR= 0.43 [0.36 0.51], p < 0.001). These results indicate that participants considered opponent status and outcome feedback in their compete decisions, independent of testosterone administration.

Importantly, testosterone modulated the effects of outcome on decisions to compete (χ^2^ (1) = 9.04, *p* = 0.003, **Figure 1C**), indicating that testosterone administration increases the relative weighting of immediate outcome feedback. Specifically, follow-up tests show that testosterone administration increased the impact of winning relative to losing on a given trial on subsequent decisions to compete. As shown in **Figure 1C**, participants who received testosterone were on average significantly more likely to compete after winning compared to losing (OR = 0.55 [0.49 0.61], p < 0.001), with the odds of competing being significantly greater than chance only after winning (Odds = 1.90 [1.25 2.88], p = 0.003), but not after losing (Odds = 1.03 [0.68 1.57], p = 0.871). This effect was not significant for placebo participants (OR = 0.85 [0.71 1.01], p = 0.125): while the odds of competing being significantly greater than chance after winning (Odds = 1.79 [1.15 2.79], p = 0.01), trend levels were still observed after losing (Odds = 1.52 [0.97 2.37], p = 0.065).

Choice latency results (Table S1, right column; **Figure 1D**) showed a main effect of Outcome (χ^2^ (1) = 12.79, p < 0.001), which was also modulated by testosterone (χ^2^ (1) = 7.71, p = 0.005). These results indicate that participants who received testosterone were not only more likely to compete again after winning (Figure 1 C), but also decided to do so relatively faster (latency difference between winning vs losing in testosterone group = 101.9, p < 0.001), while there was no such difference in the placebo group (latency difference = 12.8, p = 0.579). This indicates that feedback about winning led to relatively more impulsive decision making in individuals that received testosterone, while no such effect was observed for decision times of placebo participants.

### FMRI Results

Because we are interested in decisions to compete, we are focusing our fMRI analyses on the outcome and decision periods of the task. The outcome period is of particular interest as our behavioral analyses indicated that winning and losing on a given trial has a significant impact on subsequent behavior, particularly in participants who received testosterone.

#### Testosterone-independent activity during the outcome period

As shown **in Figure 2** (red regions, see also **Table S2**), the contrast won vs. lost during the outcome period identified a large cluster (k = 1100) in the valuation network that covers bilateral ventral striatum (VS left: -12, 11, -7; VS right: 12, 11, -7) and vmPFC (0, 56, -4), and an additional cluster in posterior parts of the temporoparietal junction (TPJ left: -39, -61, 41, k = 99; TPJ right: 45, -58, 32, k = 46). For the opposite contrast (lost vs. won, yellow regions) anterior mid cingulate cortex (6, 23, 32, k = 541), as well as a region in posterior superior temporal sulcus (-60, -37, 23, k = 92) was activated.

**Figure 2.**
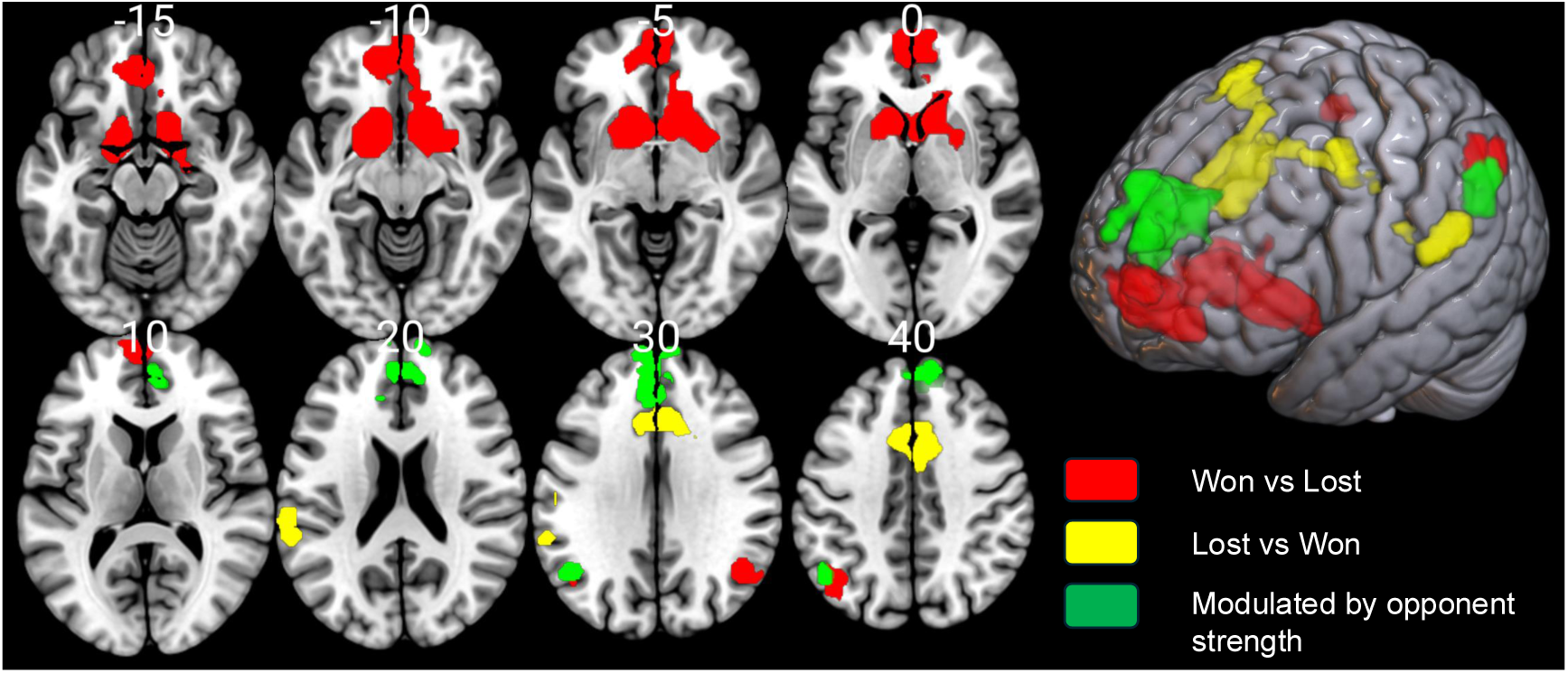
Outcome-related signals. During the outcome period, winning (vs losing, shown in red) was associated with increased activity in valuation regions in VS and vmPFC, but also social cognition regions in posterior TPJ. Losing against an opponent (vs winning, shown in yellow) was associated with activity in the mid cingulate cortex and pSTS. Finally, outcome related signal (during both win and loss) was modulated by opponent strength (shown in green) in social cognition regions including TPJ and dmPFC.

Feedback during the outcome period involved information about whether subjects won or lost against an opponent of higher, equal or lower relative ranking. Feedback therefore involves a social component, which we further investigated by testing whether activity is modulated by opponent status. We find that activity in social cognition regions that include the dorsomedial PFC (-6, 62, 26, k = 416) and posterior TPJ (-45, -58, 35, k = 87) is modulated by the status of the other player during the outcome period (green regions, **Figure 2**).

#### Testosterone modulates activity during the outcome period

Given its moderating effect on compete decisions particularly after having won or lost, it stands to reason that testosterone administration also modulates brain activity during the outcome period. To test this, we compared activations of testosterone to placebo subjects during the outcome event. **Figure 3** (see also **Table S2**) shows our results during the outcome period. Specifically, participants who received testosterone showed greater activity in FEF / posterior dlPFC (36, 5, 38, k = 54) and intraparietal sulcus (IPS: 33, -43, 41, k = 96), but these effects of testosterone are independent of whether subjects won or lost. Such significantly larger activity in regions of the canonical attention network in participants who received testosterone might reflect greater attentional orienting to stimuli that indicate relative success and social standing, which agrees well with the behavioral results and with prior theories of testosterone’s links to status-striving motivation (Eisenegger et al. 2011; Mazur and Booth 1998).

**Figure 3.**
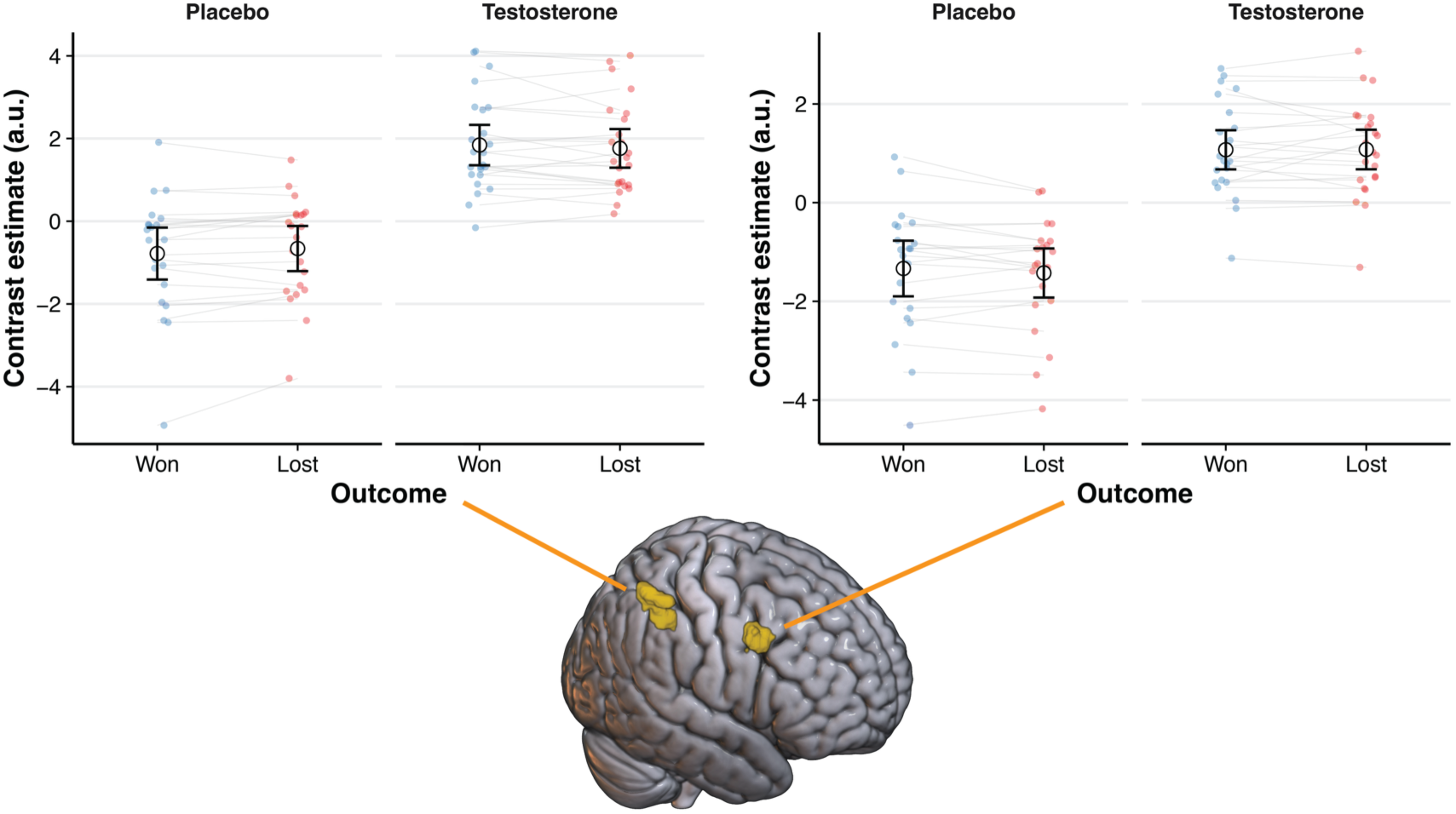
Testosterone modulates outcome-related signals. Testosterone administration increased average activation in interparietal sulcus (left) and a region including FEF / posterior region of dlPFC (right), independent of outcome.

#### Testosterone does not modulate choice-related activity

During the choice period, independent of whether subjects decided to compete or not, we observed activity in mid cingulate cortex (6, 20, 41, k = 87) and cerebellum (21, -76, -19, k = 49). A marginally non-significant effect of opponent status on choice-related activity in inferior anterior insula (-42, 20, -16, k = 38, p = 0.097, **Table S3**) was obtained when participants decided to compete. The above results apply to all participants, as we did not find any evidence that choice-related activity was modulated by testosterone.

#### Brain-Behavior relationships between outcome-related activity in valuation and social cognition regions and decisions to compete

To identify to what degree neural activity during the outcome period modulates decisions to compete, we conducted prediction analyses (Berkman and Falk 2013; Engelmann et al. 2015; Knutson et al. 2007; Knutson and Genevsky 2018). We extracted activity from those valuation (VS and vmPFC), and social cognition regions (TPJ, pSTS, dmPFC) that we observed to be active during the outcome period (see methods) and entered this information into new regression models with the goal to identify trial-by-trial influences of regional signal on decisions to compete. First, we tested whether average activity during the outcome period predicts decisions. We did not find consistent main effects indicating that regional signal predicts trial-by-trial choices on average, with the possible exception of vmPFC that showed a marginally non-significant effect (**Table S4**, all regions). Next, we inspected whether the relationship between regional neural signal and behavior was modulated by a number of key variables, including whether subjects won or lost on a given trial, the ranking of the opponent and testosterone administration. We assessed these higher-order interaction effects in separate analyses, one focusing on outcome effects and one focusing on status effects, to reduce model complexity and avoid overfitting.

#### The influence of outcome-related activity in valuation and social cognition regions on decisions to compete depends jointly on whether subjects won vs. lost and testosterone administration

Across the prediction analyses reported below we find a recurring pattern in valuation (VS, vmPFC) and social cognition (TPJ) regions. Specifically, we find that trial-by-trial neural signal during the outcome period predicts subsequent decisions to compete, but predominantly in participants who received testosterone, and in a manner that depends on whether they won or lost. The key difference across regions lies in *how* outcome modulates this relationship — VS activity amplifies the tendency to re-engage after winning, whereas TPJ activity predicts renewed competition after losing. We describe each region in turn.

First, we inspected whether outcome-related activity in valuation regions is associated with subsequent decisions (**Table S4**). We find that average activity in vmPFC during the outcome period positively predicts subsequent decisions to compete across all subjects (OR = 1.28 [1.01 1.63], p = 0.043, **Table S4** all regions). Specifically, each increase in vmPFC signal by one SD unit increases the odds of competing by 28%.

Moreover, we test whether the ability of regional signal to predict choice is jointly modulated by whether subjects won or lost on a given trial, and testosterone administration. We find such interactions in left and right ventral striatum, vmPFC and left TPJ (**Figure 4**). Activity in left ventral striatum differentially predicts decisions to compete after subjects won compared to after they lost (χ^2^ (1) = 6.81, p = 0.009). Importantly, a three-way interaction between ventral striatal signal, outcome and testosterone administration (χ^2^ (1) = 8.857, p = 0.003) indicates that the influence of ventral striatum activity on compete choices after winning compared to losing differs as a function of testosterone administration. While no effect is observed in the placebo group (interaction contrast between Outcome and VS signal: OR = 1.14 [0.43 3.05], p = 0.79), a significant modulation of compete decisions is found in the testosterone group (OR = 0.134 [0.05 0.37], p < 0.001). Specifically, participants receiving testosterone showed a 2.64-fold increase in the odds of choosing to compete after winning when VS activity was higher (OR = 2.64 [1.30 5.37], p = 0.007), and were less likely to compete after losing when VS activity was higher (OR = 0.355 [0.17 0.74], p = 0.005).

**Figure 4.**
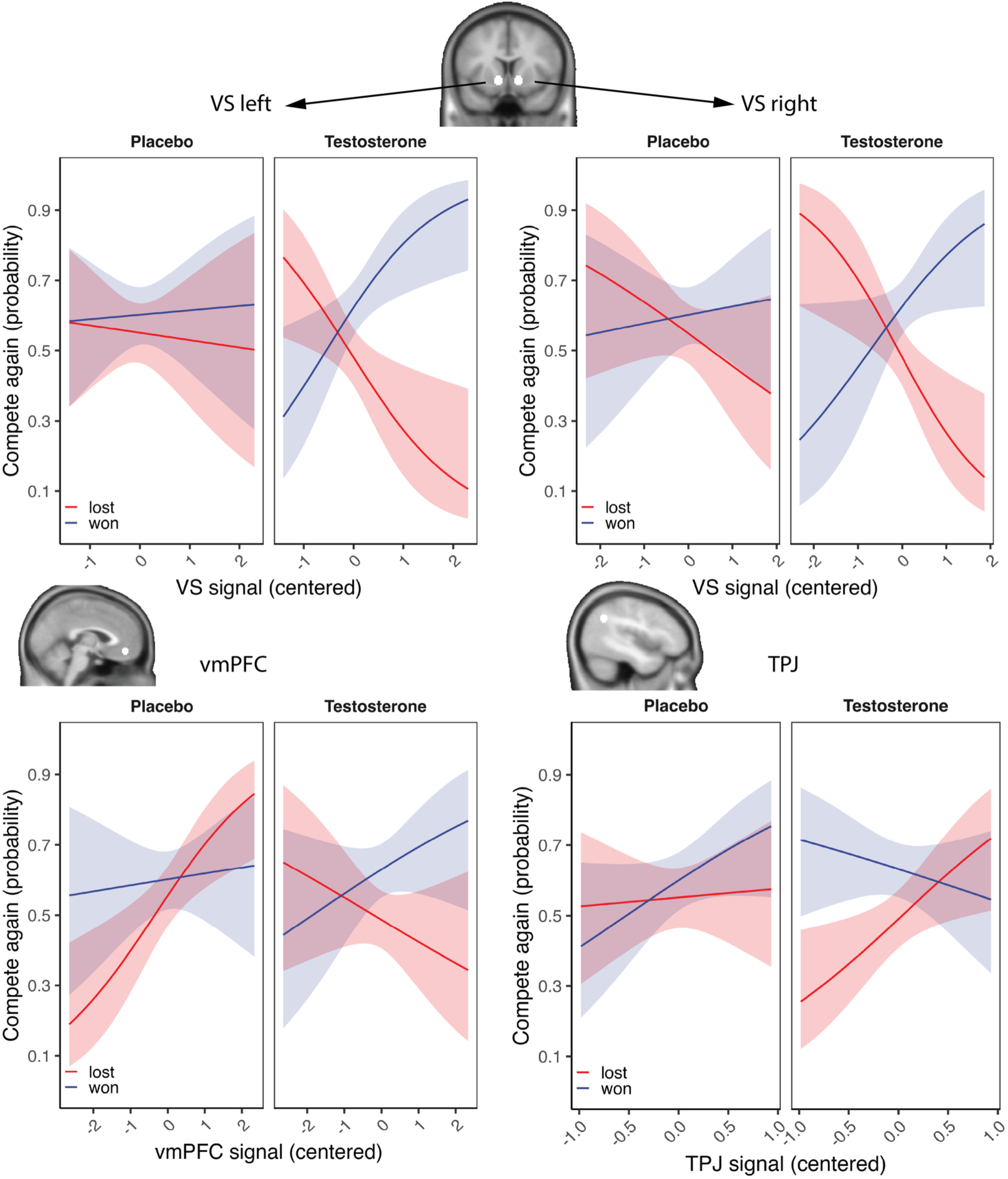
Choices to compete are influenced jointly by testosterone administration and activity in VS, vmPFC and TPJ during the outcome period. Activity in these regions is not associated with compete choices in the control group, but shows significant association with compete decisions in the Testosterone group. Regression plots reflect model estimates based on GLME logistic regressions with compete decisions as dependent variable and regional signal during the outcome period that preceded choice by 4 seconds as predictor variable.

Signal in the right ventral striatum shows a similar pattern as the left ventral striatum, but the three-way interaction did not reach significance (χ^2^ (1) = 2.55, p = 0.11). Together with the significant interaction between signal and outcome (χ^2^ (1) = 10.049, p = 0.002), these results indicate that right VS signal predicts decisions to compete as a function of outcome similarly across groups. Follow-up tests independent of testosterone administration show a decreased likelihood to compete after losing when signal in VS is low (OR = 0.52 [0.32 0.83], p = 0.007), and trend effects after winning (OR = 1.54 [0.95 2.50], p = 0.079). Due to the similar activation patterns depicted in left and right ventral striatum (**Figure 4**) and the trend level three-way interaction (p = 0.11), we conducted exploratory follow-up tests in which we probed the predictive power of right VS outcome-dependent activity for each group separately. Like in the left VS, we find a significant modulation of compete decisions in the testosterone group (interaction contrast between Outcome and VS signal: OR = 0.19 [0.07 0.51], p < 0.001), but not in the placebo group (OR = 0.58 [0.23 1.49], p = 0.26). Specifically, participants receiving testosterone showed a 2.11-fold increase in the odds of choosing to compete after winning when VS activity was higher (OR = 2.11 [1.04 4.25], p = 0.037), and were less likely to compete after losing when VS activity was higher (OR = 0.40 [0.20 0.81], p = 0.010).

Signal in the left TPJ showed a three-way interaction with outcome and testosterone administration (χ^2^ (1) = 6.43, p = 0.011). However, the pattern is opposite to that observed in the VS: while no effect is observed in the placebo group (interaction contrast between Outcome and TPJ signal: OR = 0.364 [0.10 1.40], p = 0.141), a significant modulation of compete decisions is found in the testosterone group (interaction contrast between outcome and TPJ signal: OR = 3.97 [1.13 13.92], p = 0.031). Specifically, participants who received testosterone were more likely to compete after losing but only if TPJ activity is high (OR = 2.63 [1.08 6.37], p = 0.032), while choice was not significantly predicted by TPJ activity after winning (OR = 0.66 [0.27 1.62], p = 0.366). These results indicate that participants who received testosterone were willing to compete after losing when activity in the TPJ was high, and conversely less likely to compete when TPJ activity was low.

Signal in vmPFC also interacted significantly with outcome and testosterone administration in predicting subsequent choices to compete (χ^2^ (1) = 5.86, p = 0.02). Follow-up tests show that vmPFC signal interacted with Outcome in the placebo group, but reaching only trend-level significance (OR = 1.85 [0.99 3.47], p = 0.055), but not in the testosterone group (OR = 0.60 [0.30 1.19], p = 0.141). In the placebo group, this interaction was driven by a significant increase in the odds of competing after losing when vmPFC activity was high (OR = 2.06 [1.33 3.19], p = 0.001), whereas vmPFC activity after winning did not predict subsequent choice (OR = 1.11 [0.71 1.75], p = 0.65).

#### The influence of outcome-related TPJ activity on choices to compete depends on opponent status and is independent of testosterone administration

Next, we probed the trial-by-trial signal in social cognition regions for their relationship with decisions (**Table S5**). We find that outcome-related signal from the left TPJ (-45, -58, 35) significantly predicts choice and it does so in two ways. Specifically, there is a main effect of signal (χ^2^ (1) = 10.199, p < 0.001) indicating that greater TPJ activity is associated with a higher likelihood to compete. Moreover, this effect is modulated by the status of the opponent (χ^2^ (2) = 9.55, p = 0.008). As shown in **Figure 5**, this effect indicates that specifically for the high-status opponent, TPJ signal during the outcome period is predictive of subsequent choices to compete (OR = 4.48 [2.24 8.95], p < 0.001), but not for low (OR = 1.194 [0.57 2.49], p = 0.635) and medium status opponents (OR = 1.237 [0.69 2.22], p = 0.475). The predictive effect (slope) of TPJ signal for compete decisions does not differ for control and testosterone participants across opponent status (Predictive effect for TPJ signal at Low: OR = 1.789 [0.41 7.75], p = 0.437; Medium: OR = 0.592 [0.18 1.90], p = 0.379, High: OR = 1.055 [0.27 4.19], p = 0.940), indicating that this effect is not influenced by testosterone administration. Moreover, no other social cognition region showed similar effects of opponent status.

**Figure 5.**
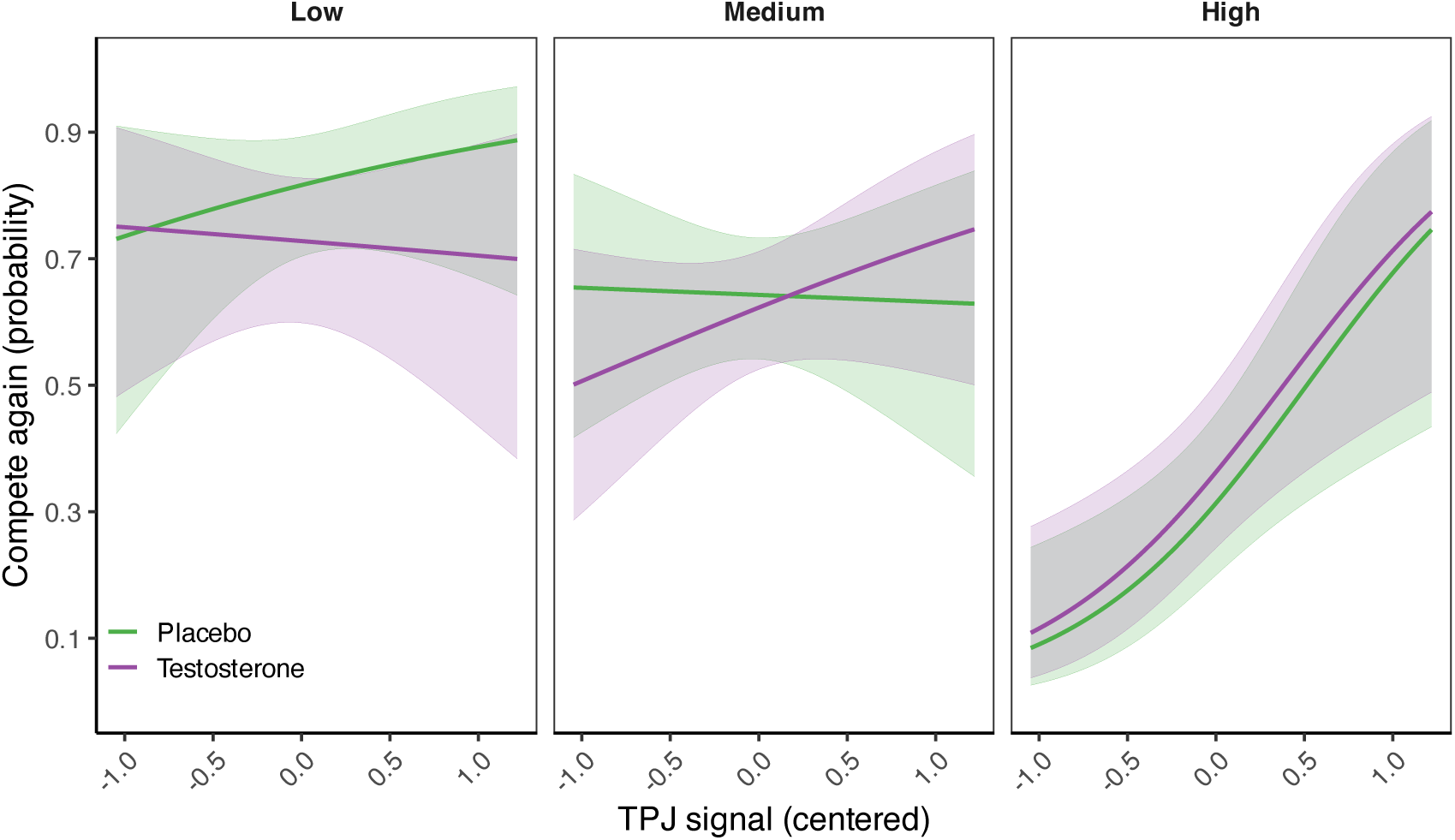
The influence of outcome-related TPJ activity on choices to compete depends on opponent status. As shown in the figure, specifically for the high-status opponent TPJ signal during the outcome period is predictive of subsequent choices to compete.

## Discussion

We investigated the behavioral and neurobiological effect of testosterone administration on decisions to compete after winning or losing against opponents of varying status. Two main findings emerge from this study: First, participants generally adjusted decisions based on status and feedback, but those given testosterone showed greater sensitivity to outcomes: they were more likely to re-compete after wins, less likely after losses, and made decisions faster. Second, trial-level analyses showed that activity in the left ventral striatum and TPJ predicted subsequent competition decisions only in the testosterone group. Striatal activity was associated with re-competing after wins, whereas TPJ activity predicted re-engagement after losses.

In our task, participants made decisions about whether to compete against an opponent based on two types of information: 1. they were informed about their opponent’s status, which provides participants with an average success likelihood against their opponent and 2. they received feedback about whether on a current trial they won or lost against the opponent. We find that participants in both placebo and testosterone groups strongly weighted information about opponent status in their decisions and were significantly less likely to compete against higher status than lower status individuals (the odds of competing against a low-status opponent were 7.2 times greater than competing against a high-status opponent). Strategic decisions about whether to compete with an opponent should not just reflect ability ratings from past performance, but also fluctuations reflected by feedback about the outcome of the current competition. Participants in both groups also accounted for whether they won or lost in the current competition in their decisions to compete (participants were 1.47 times more likely to compete after winning compared to losing).

Importantly, we find that testosterone administration changes how this information is used in the decision to compete against an opponent. Specifically, the weighting of immediate feedback was increased under testosterone, such that participants were about 1.9 times more likely to compete after winning, whereas after losing they were only about half as likely to compete compared to winning (OR = 0.55). This pattern of results is consistent with previous correlational evidence (Mehta et al. 2008). The finding that decisions are faster after winning in participants that received testosterone administration, but not in the placebo group, further underlines that testosterone participants made decisions more impulsively after winning. Together, these behavioral results indicate that testosterone administration caused a shift in how important information is processed in competitive decision-making, leading to an increase in the weighting of immediate information, over longer-term strategic information.

Our neural results mirror this effect. We identify a network of regions that process outcome-related information in all participants (independent of testosterone administration) in bilateral ventral striatum, vmPFC, temporoparietal junction and anterior cingulate. Interestingly, outcome related signal in temporoparietal junction and dorsomedial prefrontal cortex tracked opponent status, indicating that social cognition regions play an important role in integrating the two different pieces of information (opponent status and recent opponent success) important for subsequent decisions to compete. This agrees with the role of these regions in strategic decision contexts that has been previously demonstrated (Bitsch et al. 2018; Chang et al. 2023; Engelmann et al. 2019; Morishima et al. 2012; Rilling et al. 2004).

To directly relate the outcome related activation patterns to subsequent decisions, we conducted prediction analyses that ask to what extent trial-by-trial signal variation in the outcome network predicts compete decisions. We find that activity in left (and similarly in right) ventral striatum and TPJ predicts subsequent compete decisions, but only in participants that received testosterone administration, while vmPFC activity predicts decisions only in Placebo participants. Moreover, in participants that received testosterone, the predictive effects of striatum activity depended on whether subjects won or lost on a given trial and therefore reflects the behavioral results. Together with the behavioral results, the findings indicate that testosterone administration leads to greater responsivity to immediate feedback about opponent performance, which is moderated by regional activity in a wider network that includes reward (ventral striatum, vmPFC) and social cognition regions (TPJ).

Our result that ventral striatum activity modulates compete decisions after testosterone administration is consistent with its well-established role within the valuation and reward network. The ventral striatum processes not only primary rewards, but also social rewards, such as outperforming an opponent (Bault et al. 2011; Fareri and Delgado 2014; Fliessbach et al. 2007). Prior work has further shown that testosterone modulates activity in ventral striatum, both as a function of endogenous testosterone in adolescents (Op De Macks et al. 2011) and in a placebo-controlled testosterone administration study in women (Hermans et al. 2010). Extending these findings, our results show that increased striatal activity after winning was associated with a higher likelihood to compete, whereas increased striatal activity after losing was associated with a lower likelihood to compete – an effect that was only observed in participants who received testosterone. Rather than simply boosting reward responses after success, testosterone appears to enhance the sensitivity of the ventral striatum to the motivational meaning of competitive outcomes, driving approach after winning and withdrawal after losing. This interpretation aligns with non-human animal evidence showing that testosterone treatment coupled with winning experience enhances competitive behavior via androgen receptor sensitivity in the ventral striatum (Fuxjager et al. 2010), and extends it by suggesting that testosterone equally amplifies the behavioral impact of losing. While the amygdala has been identified as a key mediator of testosterone’s motivational approach-avoidance responses to aversive social stimuli (Radke et al. 2015), the present findings extend this by implicating the ventral striatum in outcome-driven competitive decisions — possibly reflecting the more complex motivational computation required when deciding whether to re-engage a specific opponent after a loss.

The differential role of the TPJ in competitive decisions after testosterone administration is in line with prior research indicating (1) a general involvement of the TPJ in social cognition in the context of strategic interactions (Chang et al. 2023; Engelmann et al. 2019; Rilling et al. 2004), and (2) testosterone-specific modulation of TPJ activity in related social interactive contexts (Chen et al. 2016; Ou et al. 2021). More broadly, numerous studies implicate the TPJ in social cognition (Decety and Lamm 2007; Mar 2011; Molenberghs et al. 2016; Schurz et al. 2014; Van Overwalle 2009), and, importantly, in strategic decision-making within economic games (Chang et al. 2023; Engelmann et al. 2019; Fukui et al. 2006; Ou et al. 2021; Schurz et al. 2014). Our current findings extend this work by showing that when participants decided whether to re-engage an opponent who had just defeated them, enhanced TPJ activity predicted a greater likelihood of choosing to compete, but only following testosterone administration. Such situations require strategic reasoning about whether it is possible to overcome a stronger opponent. Prior work suggests that similar choices in the context of trust games engage mentalizing processes subserved by the TPJ, enabling individuals to estimate the intentions of an opponent and the likelihood of interaction success (Chang et al. 2023; Engelmann et al. 2019). We propose that in the present context, testosterone amplifies TPJ contributions to mentalizing, thereby biasing individuals toward renewed competition despite recent defeat.

Overall, our results demonstrate that testosterone modulates brain-behavior relationships in competitive decision-making by amplifying the influence of outcome-related activity in valuation (ventral striatum) and social cognition (bilateral TPJ) regions on subsequent decisions to compete. Together with the behavioral finding that testosterone selectively enhanced willingness to compete after winning, and the increased recruitment of the canonical attention network during outcome processing, these results converge on a coherent account: testosterone increases the weighting of immediate, salient outcome information — amplifying both the neural sensitivity to competitive outcomes and their downstream influence on behavior.

## Use of AI-assisted technologies

All analyses, results, and writing were conducted by the authors; AI-assisted tools (ChatGPT, OpenAI) were used in a limited capacity to assist with R coding (e.g., extracting contrasts with *emmeans* and generating plots with *ggplot2*) and minor text editing.

## Supplemental Materials

**Table S1.**
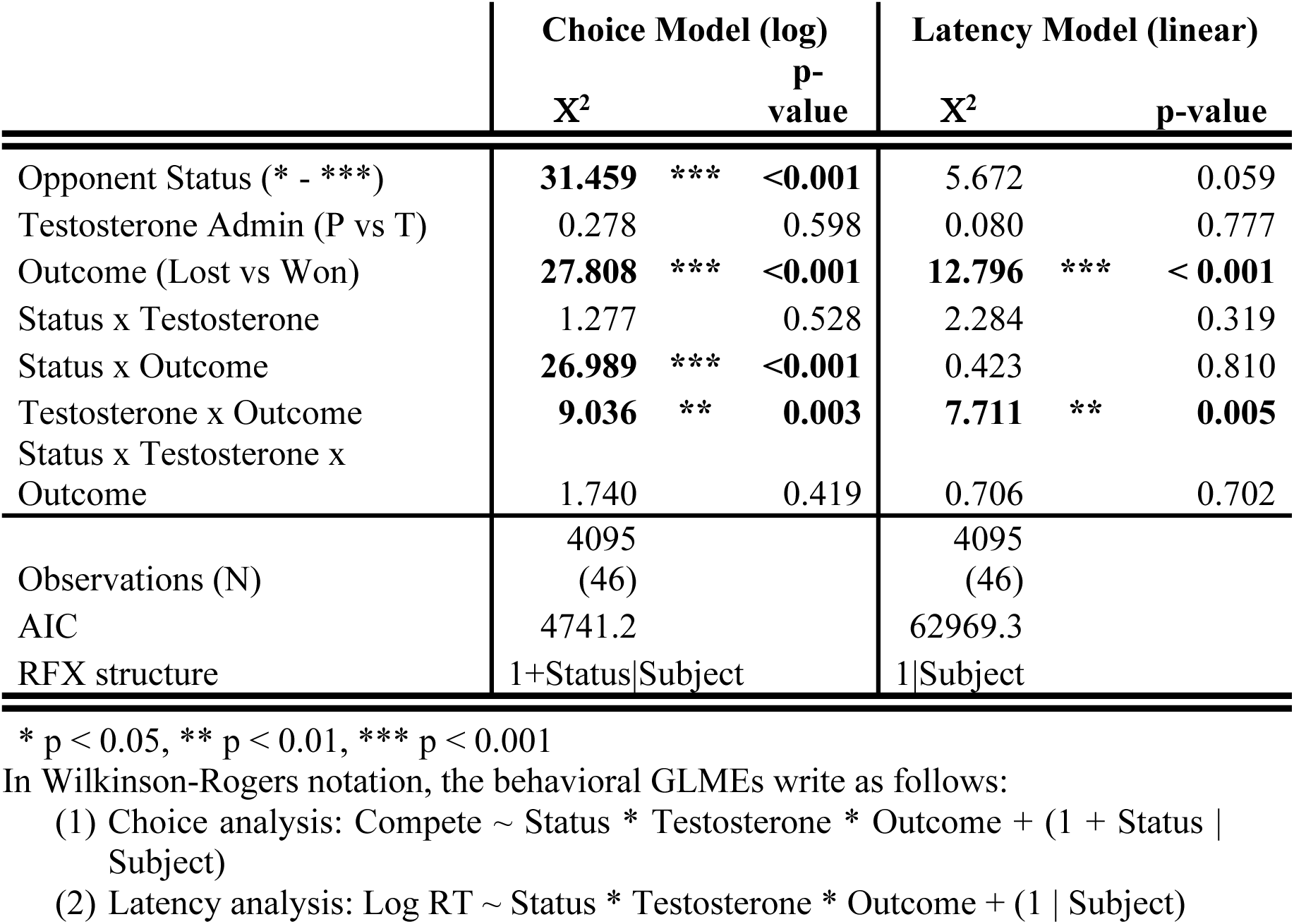
Behavioral Results. This table shows the output from behavioral analyses using a logistic model for the choice data, and a linear model for the log normalized reaction time data using Likelihood Ratio Tests (LRT) to estimate p values. Results are based on the maximal model structure that converged incorporating subject-specific random intercepts and random slopes for the within-subject factors Opponent Status. Note that response latency results generate nearly identical p-values for log normalized RT.

**Figure S1.**
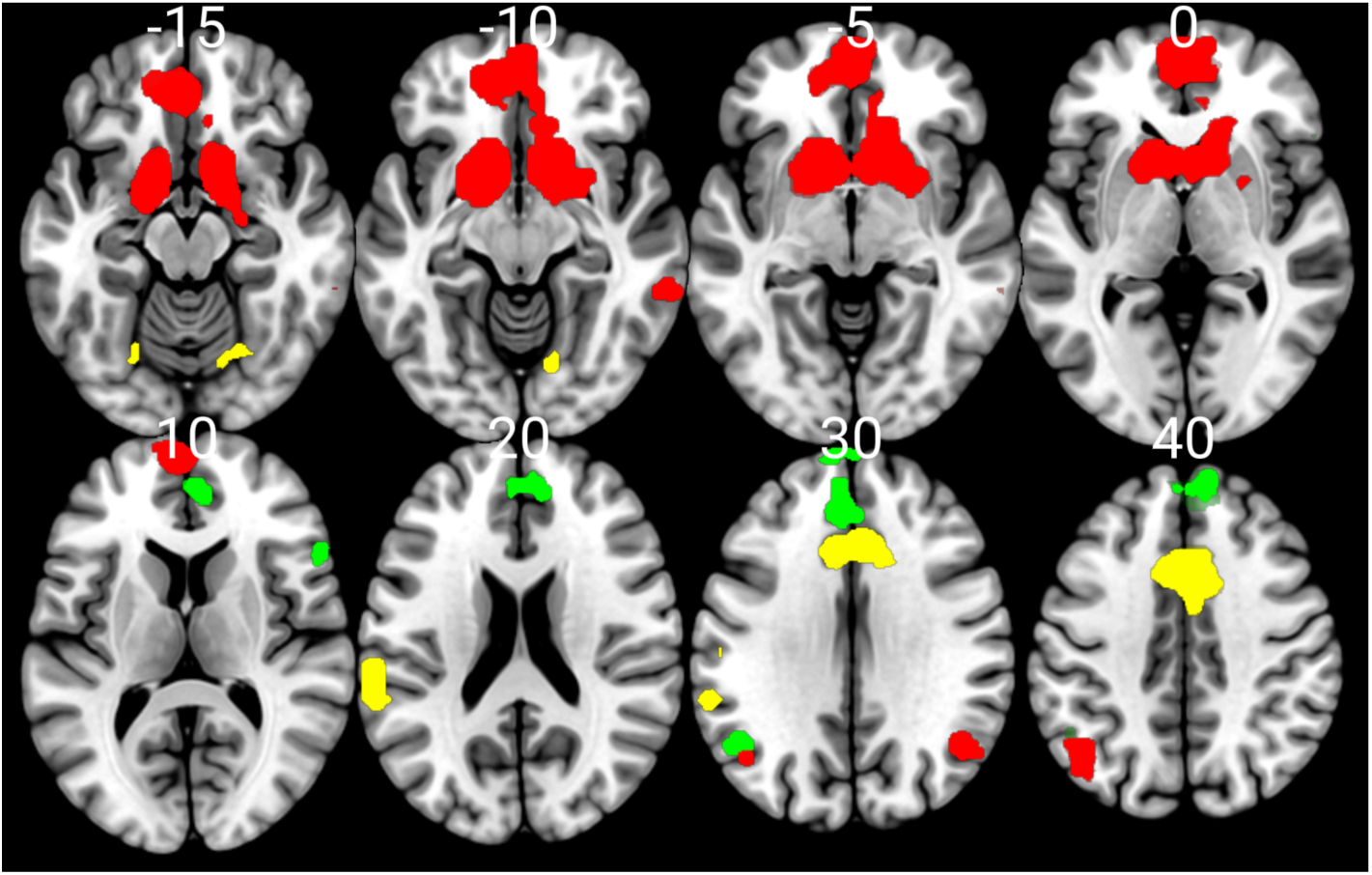
Conjunction showing consistency of main activation patterns during the outcome period over models implementing different temporal shifts (0s, +1s, +3s). Activation maps were first thresholded with a cluster forming threshold of p < 0.001 and a cluster extent of k=20 and were then entered into a *logical-and* conjunction analysis using imcalc (by multiplying all maps), to identify voxels that survive in all three maps. Red reflects the contrast Won vs Lost, Yellow reflects Lost vs Won and Green shows outcome-related activity modulated by opponent status.

**Table S2.** Activations during the outcome period (cluster-forming height threshold = p<0.001, FWE-corrected at the cluster level).

| Contra<br>st | Region | L/R | Cluster<br>size | Cluster-level<br>p(FWE-corr) | Peak<br>T | X | Y | Z |
| --- | --- | --- | --- | --- | --- | --- | --- | --- |
| *Outcome Event*<br>(*FWEc* = 229) |  |  |  |  |  |  |  |  |
|  | Occipital lobe* | bilateral | 5487 | <0.001 | 13.29 | -39 | -79 | -10 |
|  | Intraparietal Sulcus | left | 404 | <0.001 | 7.76 | -39 | -52 | 56 |
|  | Middle Frontal Gyrus | right | 335 | <0.001 | 7.7 | 48 | 35 | 29 |
|  | Putamen | left | 229 | <0.001 | 5.89 | -21 | 11 | 8 |
| *Won vs Lost*<br>(*FWEc* = 99) |  |  |  |  |  |  |  |  |
|  | Ventral Striatum, Nacc | left | 1107 | <0.001 | 10.68 | -12 | 11 | -7 |
|  | Ventral Striatum,<br>Nacc* | right |  |  | 9.55 | 12 | 11 | -7 |
|  | Anterior vmPFC* | bilateral |  |  | 6.12 | 0 | 56 | -4 |
|  | Temporoparietal<br>Junction | left | 99 | 0.002 | 5.06 | -39 | -61 | 41 |
|  | *Temporoparietal*<br>*Junction* | right | 46 | 0.059 | 4.71 | 45 | -58 | 32 |
|  | *dorsolateral PFC* | right | 43 | 0.074 | 4.12 | 24 | 29 | 47 |
| *Lost vs. Won*<br>(*FWEc* = 92) |  |  |  |  |  |  |  |  |
|  | Anterior Cingulate<br>Cortex | bilateral | 541 | <0.001 | 7.02 | 6 | 23 | 32 |
|  | Posterior STS | left | 92 | 0.003 | 5.04 | -60 | -37 | 23 |
| *Won vs. Lost x Opponent*<br>*Status*<br>(*FWEc* = 87) |  |  |  |  |  |  |  |  |
|  | dorsomedial PFC | bilateral | 416 | <0.001 | 5.68 | -6 | 62 | 26 |
|  | Temporoparietal<br>Junction | left | 87 | 0.003 | 4.93 | -45 | -58 | 35 |
| *Outcome: Testosterone >*<br>*Placebo*<br>(*FWEc* = 54) |  |  |  |  |  |  |  |  |
|  | FEF / posterior dlPFC | right | 54 | 0.036 | 5.34 | 36 | 5 | 38 |
|  | Intraparietal Sulcus | right | 96 | 0.002 | 4.9 | 33 | -43 | 41 |

\* indicates subcluster within larger activation cluster, italics indicate regions that are marginally non-significant.

**Table S3.**
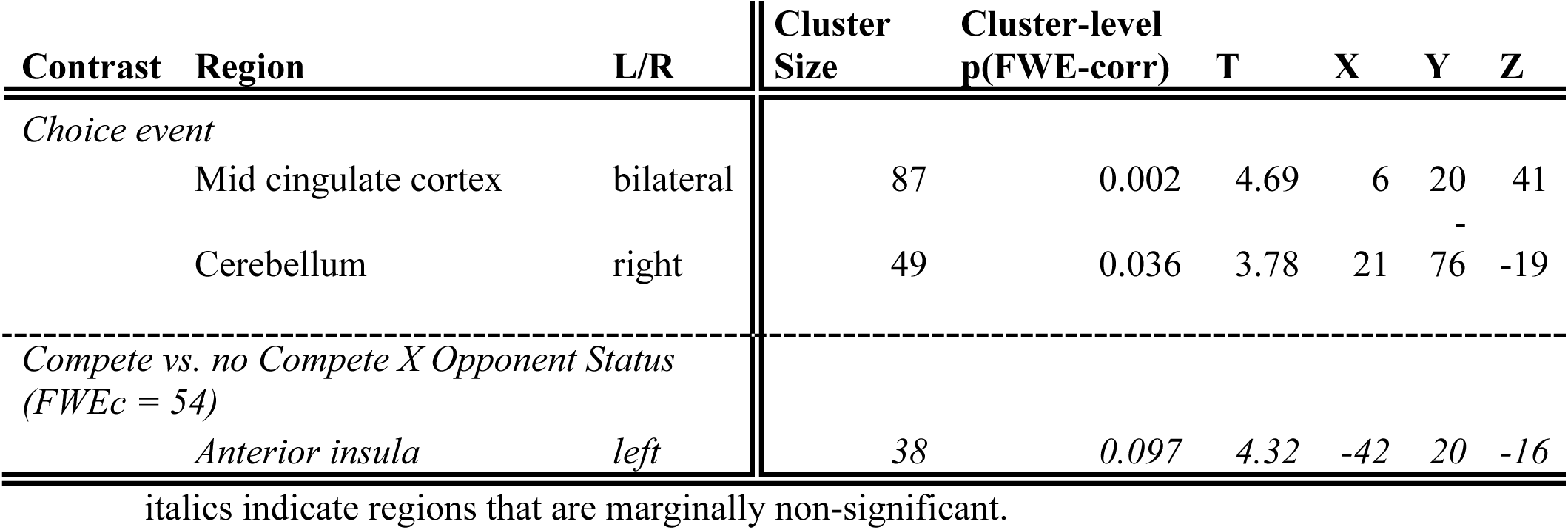
Activations during the choice period (cluster-forming height threshold = p<0.001, FWE-corrected at the cluster level).

**Table S4.** Prediction analyses of outcome-related activity focusing on outcome on the current trial. VS, vmPFC and TPJ activity are predictive of compete decisions under specific conditions.

|  | All regions | VS left | VS right | vmPFC | ACC | TPJ left | TPJ right |
| --- | --- | --- | --- | --- | --- | --- | --- |
| VS Left | 0.001 | 0.028 |  |  |  |  |  |
| VS right | 0.886 |  | 0.426 |  |  |  |  |
| vmPFC | **4.086** * |  |  | 2.520 |  |  |  |
| ACC | 0.075 |  |  |  | 0.034 |  |  |
| TPJ left | 2.749 + |  |  |  |  | 2.579 |  |
| TPJ right | 0.650 |  |  |  |  |  | 0.011 |
| Signal X Outcome |  | **6.801** ** | **10.049** ** | 0.049 | 1.982 | 0.152 | 0.085 |
| Signal X Testosterone |  | 0.000 | 0.040 | 3.522 + | 2.168 | 0.186 | 0.127 |
| Signal x Outcome X Test |  | **8.857** ** | 2.552 | **5.654** * | 0.703 | **6.433** * | 1.842 |
| Outcome | 31.526 *** | 32.329 *** | 33.607 *** | 32.316 *** | 33.352 *** | 32.538 *** | 33.617 *** |
| Testosterone Administration | 0.003 | 0.015 | 0.001 | 0.250 | 0.002 | 0.013 | 0.000 |
| Status | 31.357 *** | 31.488 *** | 31.254 *** | 35.874 *** | 30.925 *** | 31.677 *** | 30.967 *** |
| Outcome X Testosterone | 7.140 ** | 6.475 * | 6.440 * | 7.838 ** | 6.192 * | 7.118 ** | 7.139 ** |
| Observations (N) | 4086 (46) | 4086 (46) | 4086 (46) | 4086 (46) | 4086 (46) | 4086 (46) | 4086 (46) |
| AIC | 4748.5 | 4736.4 | 4739 | 4747.7 | 4747 | 4742.6 | 4749.9 |
| RFX structure | 1+Status\|Subject | 1+Status\|Subject | 1+Status\|Subject | 1+Status\|Subject | 1+Status\|Subject | 1+Status\|Subject | 1+Status\|Subject |

\* $p < 0.05$ , \*\* $p < 0.01$ , \*\*\* $p < 0.001$
Outcome-related neural prediction analyses in Wilkinson-Rogers notation:
Compete ~ VS<sub>l</sub>\_signal + VS<sub>r</sub>\_signal + vmPFC\_signal + ACC\_signal + TPJ<sub>l</sub>\_signal + TPJ<sub>r</sub>\_signal + outcome \* treatment + status + (1 + Status |Subject)

**Table S5.** Prediction analyses of outcome-related activity focusing on opponent status.

|  | All regions | TPJ left | TPJ left | TPJ right | dmPFC |
| --- | --- | --- | --- | --- | --- |
|  | Chisq | Chisq | Chisq | Chisq | Chisq |
| TPJ left | 0.297 | 2.131 |  |  |  |
| TPJ left | 10.199 *** |  | 10.025 ** |  |  |
| TPJ right | 3.202 + |  |  | 0.101 |  |
| dmPFC | 0.749 |  |  |  | 1.007 |
| Signal X Status |  | 3.748 | 9.550 ** | 4.129 | 1.094 |
| Signal x Testosterone |  | 0.000 | 0.009 | 0.110 | 1.622 |
| Signal X Status X Test |  | 0.869 | 1.365 | 0.399 | 3.043 |
| Status | 31.999 *** | 36.330 *** | 37.219 *** | 30.637 *** | 31.827 *** |
| Testosterone | 0.227 | 0.261 | 0.193 | 0.252 | 0.219 |
| Outcome | 34.144 *** | 35.399 *** | 31.808 *** | 36.520 *** | 36.718 *** |
| Status x Testosterone | 1.387 | 1.295 | 1.398 | 1.269 | 1.388 |
| Observations | 4086 (46) | 4086 (46) | 4086 (46) | 4086 (46) | 4086 (46) |
| AIC | 4745.2 | 4742.3 | 4752.1 | 4759.1 | 4757.1 |
| RFX structure | 1+Status\|Subject | 1+Status\|\|Subject | 1+Status\|\|Subject | 1+Status\|Subject | 1+Status\|Subject |

\* $p < 0.05$ , \*\* $p < 0.01$ , \*\*\* $p < 0.001$
Status-related neural prediction analyses in Wilkinson-Rogers notation:
Compete ~ VS<sub>l</sub>\_signal + VS<sub>r</sub>\_signal + vmPFC\_signal + ACC\_signal + TPJ<sub>l</sub>\_signal + TPJ<sub>r</sub>\_signal + status \* treatment + outcome + (1 + Status |Subject)

## References

Barr, Dale J. 2013. “Random Effects Structure for Testing Interactions in Linear Mixed-Effects Models.” Frontiers in Psychology 4. doi:10.3389/fpsyg.2013.00328.

Bartra, Oscar, Joseph T. McGuire, and Joseph W. Kable. 2013. “The Valuation System: A Coordinate-Based Meta-Analysis of BOLD fMRI Experiments Examining Neural Correlates of Subjective Value.” NeuroImage 76:412–27. doi:10.1016/j.neuroimage.2013.02.063.

Bates, Douglas, Martin Mächler, Ben Bolker, and Steve Walker. 2015. “Fitting Linear Mixed-Effects Models Using Lme4.” Journal of Statistical Software 67:1–48. doi:10.18637/jss.v067.i01.

Bault, Nadège, Mateus Joffily, Aldo Rustichini, and Giorgio Coricelli. 2011. “Medial Prefrontal Cortex and Striatum Mediate the Influence of Social Comparison on the Decision Process.” Proceedings of the National Academy of Sciences 108(38):16044–49. doi:10.1073/pnas.1100892108.

Beehner, J. C., T. J. Bergman, D. L. Cheney, R. M. Seyfarth, and P. L. Whitten. 2006. “Testosterone Predicts Future Dominance Rank and Mating Activity among Male Chacma Baboons.” Behavioral Ecology and Sociobiology 59(4):469–79. doi:10.1007/s00265-005-0071-2.

Berkman, Elliot T., and Emily B. Falk. 2013. “Beyond Brain Mapping: Using Neural Measures to Predict Real-World Outcomes.” Current Directions in Psychological Science 22(1):45–50. doi:10.1177/0963721412469394.

Bitsch, Florian, Philipp Berger, Arne Nagels, Irina Falkenberg, and Benjamin Straube. 2018. “The Role of the Right Temporo–Parietal Junction in Social Decision-Making.” Human Brain Mapping 39(7):3072–85. doi:10.1002/hbm.24061.

Bolker, Benjamin M., Mollie E. Brooks, Connie J. Clark, Shane W. Geange, John R. Poulsen, M. Henry H. Stevens, and Jada-Simone S. White. 2009. “Generalized Linear Mixed Models: A Practical Guide for Ecology and Evolution.” Trends in Ecology & Evolution 24(3):127–35. doi:10.1016/j.tree.2008.10.008.

Bos, Peter A., Erno J. Hermans, Nick F. Ramsey, and Jack Van Honk. 2012. “The Neural Mechanisms by Which Testosterone Acts on Interpersonal Trust.” NeuroImage 61(3):730–37. doi:10.1016/j.neuroimage.2012.04.002.

Bos, Peter A., Jack Van Honk, Nick F. Ramsey, Dan J. Stein, and Erno J. Hermans. 2013. “Testosterone Administration in Women Increases Amygdala Responses to Fearful and Happy Faces.” Psychoneuroendocrinology 38(6):808–17. doi:10.1016/j.psyneuen.2012.09.005.

Carter, R. McKell, and Scott A. Huettel. 2013. “A Nexus Model of the Temporal–Parietal Junction.” Trends in Cognitive Sciences 17(7):328–36. doi:10.1016/j.tics.2013.05.007.

Cashdan, Elizabeth. 1995. “Hormones, Sex, and Status in Women.” Hormones and Behavior 29(3):354–66. doi:10.1006/hbeh.1995.1025.

Chang, Li-Ang, Konstantinos Armaos, Lotte Warns, Ava Q. Ma de Sousa, Femke Paauwe, Christin Scholz, and Jan B. Engelmann. 2023. “Mentalizing in an Economic Games Context Is Associated with Enhanced Activation and Connectivity in the Left Temporoparietal Junction.” Social Cognitive and Affective Neuroscience 18(1):nsad023. doi:10.1093/scan/nsad023.

Chen, Chenyi, Jean Decety, Pin-Chia Huang, Chin-Yau Chen, and Yawei Cheng. 2016. “Testosterone Administration in Females Modulates Moral Judgment and Patterns of Brain Activation and Functional Connectivity.” Human Brain Mapping 37(10):3417–30. doi:10.1002/hbm.23249.

Decety, Jean, and Claus Lamm. 2007. “The Role of the Right Temporoparietal Junction in Social Interaction: How Low-Level Computational Processes Contribute to Meta-Cognition.” The Neuroscientist 13(6):580–93. doi:10.1177/1073858407304654.

Desmond, John E., and Gary H. Glover. 2002. “Estimating Sample Size in Functional MRI (fMRI) Neuroimaging Studies: Statistical Power Analyses.” Journal of Neuroscience Methods 118(2):115–28. doi:10.1016/S0165-0270(02)00121-8.

Dreher, Jean-Claude, Simon Dunne, Agnieszka Pazderska, Thomas Frodl, John J. Nolan, and John P. O’Doherty. 2016. “Testosterone Causes Both Prosocial and Antisocial Status-Enhancing Behaviors in Human Males.” Proceedings of the National Academy of Sciences 113(41):11633–38. doi:10.1073/pnas.1608085113.

Eisenegger, C., M. Naef, R. Snozzi, M. Heinrichs, and E. Fehr. 2010. “Prejudice and Truth about the Effect of Testosterone on Human Bargaining Behaviour.” Nature 463(7279):356–59. doi:10.1038/nature08711.

Eisenegger, Christoph, Johannes Haushofer, and Ernst Fehr. 2011. “The Role of Testosterone in Social Interaction.” Trends in Cognitive Sciences 15(6):263–71. doi:10.1016/j.tics.2011.04.008.

Engelmann, Jan B., Friederike Meyer, Ernst Fehr, and Christian C. Ruff. 2015. “Anticipatory Anxiety Disrupts Neural Valuation during Risky Choice.” Journal of Neuroscience 35(7):3085–99. doi:10.1523/JNEUROSCI.2880-14.2015.

Engelmann, Jan B., Friederike Meyer, Christian C. Ruff, and Ernst Fehr. 2019. “The Neural Circuitry of Affect-Induced Distortions of Trust.” Science Advances 5(3):eaau3413. doi:10.1126/sciadv.aau3413.

Fareri, Dominic S., and Mauricio R. Delgado. 2014. “Differential Reward Responses during Competition against In- and out-of-Network Others.” Social Cognitive and Affective Neuroscience 9(4):412–20. doi:10.1093/scan/nst006.

Fliessbach, K., B. Weber, P. Trautner, T. Dohmen, U. Sunde, C. E. Elger, and A. Falk. 2007. “Social Comparison Affects Reward-Related Brain Activity in the Human Ventral Striatum.” Science 318(5854):1305–8. doi:10.1126/science.1145876.

Fukui, Hiroki, Toshiya Murai, Jun Shinozaki, Toshihiko Aso, Hidenao Fukuyama, Takuji Hayashi, and Takashi Hanakawa. 2006. “The Neural Basis of Social Tactics: An fMRI Study.” NeuroImage 32(2):913–20. doi:10.1016/j.neuroimage.2006.03.039.

Fuxjager, Matthew J., Robin M. Forbes-Lorman, Dylan J. Coss, Catherine J. Auger, Anthony P. Auger, and Catherine A. Marler. 2010. “Winning Territorial Disputes Selectively Enhances Androgen Sensitivity in Neural Pathways Related to Motivation and Social Aggression.” Proceedings of the National Academy of Sciences 107(27):12393–98. doi:10.1073/pnas.1001394107.

Grinband, Jack, Tor D. Wager, Martin Lindquist, Vincent P. Ferrera, and Joy Hirsch. 2008. “Detection of Time-Varying Signals in Event-Related fMRI Designs.” NeuroImage 43(3):509–20. doi:10.1016/j.neuroimage.2008.07.065.

Heany, Sarah J., Jack van Honk, Dan J. Stein, and Samantha J. Brooks. 2016. “A Quantitative and Qualitative Review of the Effects of Testosterone on the Function and Structure of the Human Social-Emotional Brain.” Metabolic Brain Disease 31(1):157–67. doi:10.1007/s11011-015-9692-y.

Hermans, Erno J., Peter A. Bos, Lindsey Ossewaarde, Nick F. Ramsey, Guillén Fernández, and Jack Van Honk. 2010. “Effects of Exogenous Testosterone on the Ventral Striatal BOLD Response during Reward Anticipation in Healthy Women.” NeuroImage 52(1):277–83. doi:10.1016/j.neuroimage.2010.04.019.

Huettel, Scott A., and Gregory McCarthy. 2001. “The Effects of Single-Trial Averaging upon the Spatial Extent of fMRI Activation.” NeuroReport 12(11):2411.

Hutschemaekers, Moniek H. M., Rianne A. de Kleine, Gert-Jan Hendriks, Mirjam Kampman, and Karin Roelofs. 2021. “The Enhancing Effects of Testosterone in Exposure Treatment for Social Anxiety Disorder: A Randomized Proof-of-Concept Trial.” Translational Psychiatry 11(1):432. doi:10.1038/s41398-021-01556-8.

Hutschemaekers, Moniek H. M., Rianne A. de Kleine, Mirjam Kampman, Jasper A. J. Smits, and Karin Roelofs. 2023. “Social Avoidance and Testosterone Enhanced Exposure Efficacy in Women with Social Anxiety Disorder: A Pilot Investigation.” Psychoneuroendocrinology 158:106372. doi:10.1016/j.psyneuen.2023.106372.

Knutson, Brian, and Alexander Genevsky. 2018. “Neuroforecasting Aggregate Choice.” Current Directions in Psychological Science 27(2):110–15. doi:10.1177/0963721417737877.

Knutson, Brian, and Stephanie M. Greer. 2008. “Anticipatory Affect: Neural Correlates and Consequences for Choice.” Philosophical Transactions of the Royal Society B: Biological Sciences 363(1511):3771–86. doi:10.1098/rstb.2008.0155.

Knutson, Brian, Scott Rick, G. Elliott Wimmer, Drazen Prelec, and George Loewenstein. 2007. “Neural Predictors of Purchases.” Neuron 53(1):147–56. doi:10.1016/j.neuron.2006.11.010.

Lenth, Russell V. 2016. “Least-Squares Means: The R Package Lsmeans.” Journal of Statistical Software 69:1–33. doi:10.18637/jss.v069.i01.

Losecaat Vermeer, A. B., I. Krol, C. Gausterer, B. Wagner, C. Eisenegger, and C. Lamm. 2020. “Exogenous Testosterone Increases Status-Seeking Motivation in Men with Unstable Low Social Status.” Psychoneuroendocrinology 113:104552. doi:10.1016/j.psyneuen.2019.104552.

Mar, Raymond A. 2011. “The Neural Bases of Social Cognition and Story Comprehension.” Annual Review of Psychology 62(1):103–34. doi:10.1146/annurev-psych-120709-145406.

Matuschek, Hannes, Reinhold Kliegl, Shravan Vasishth, Harald Baayen, and Douglas Bates. 2017. “Balancing Type I Error and Power in Linear Mixed Models.” Journal of Memory and Language 94:305–15. doi:10.1016/j.jml.2017.01.001.

Mazur, Allan, and Allan Booth. 1998. “Testosterone and Dominance in Men.” Behavioral and Brain Sciences 21(3):363–363. doi:10.1017/S0140525X98221224.

Mehta, Pranjal H., Amanda C. Jones, and Robert A. Josephs. 2008. “The Social Endocrinology of Dominance: Basal Testosterone Predicts Cortisol Changes and Behavior Following Victory and Defeat.” Journal of Personality and Social Psychology 94(6):1078–93. doi:10.1037/0022-3514.94.6.1078.

Mehta, Pranjal H., Veerle van Son, Keith M. Welker, Smrithi Prasad, Alan G. Sanfey, Ale Smidts, and Karin Roelofs. 2015. “Exogenous Testosterone in Women Enhances and Inhibits Competitive Decision-Making Depending on Victory–Defeat Experience and Trait Dominance.” Psychoneuroendocrinology 60:224–36. doi:10.1016/j.psyneuen.2015.07.004.

Molenberghs, Pascal, Halle Johnson, Julie D. Henry, and Jason B. Mattingley. 2016. “Understanding the Minds of Others: A Neuroimaging Meta-Analysis.” Neuroscience & Biobehavioral Reviews 65:276–91. doi:10.1016/j.neubiorev.2016.03.020.

Morishima, Yosuke, Daniel Schunk, Adrian Bruhin, Christian C. Ruff, and Ernst Fehr. 2012. “Linking Brain Structure and Activation in Temporoparietal Junction to Explain the Neurobiology of Human Altruism.” Neuron 75(1):73–79. doi:10.1016/j.neuron.2012.05.021.

Muscatell, Keely A., Sylvia A. Morelli, Emily B. Falk, Baldwin M. Way, Jennifer H. Pfeifer, Adam D. Galinsky, Matthew D. Lieberman, Mirella Dapretto, and Naomi I. Eisenberger. 2012. “Social Status Modulates Neural Activity in the Mentalizing Network.” NeuroImage 60(3):1771–77. doi:10.1016/j.neuroimage.2012.01.080.

Nave, G., A. Nadler, D. Dubois, D. Zava, C. Camerer, and H. Plassmann. 2018. “Single-Dose Testosterone Administration Increases Men’s Preference for Status Goods.” Nature Communications 9(1):2433. doi:10.1038/s41467-018-04923-0.

Op De Macks, Zdeňa A., Bregtje Gunther Moor, Sandy Overgaauw, Berna Güroğlu, Ronald E. Dahl, and Eveline A. Crone. 2011. “Testosterone Levels Correspond with Increased Ventral Striatum Activation in Response to Monetary Rewards in Adolescents.” Developmental Cognitive Neuroscience 1(4):506–16. doi:10.1016/j.dcn.2011.06.003.

Ou, Jianxin, Yin Wu, Yang Hu, Xiaoxue Gao, Hong Li, and Philippe N. Tobler. 2021. “Testosterone Reduces Generosity through Cortical and Subcortical Mechanisms.” Proceedings of the National Academy of Sciences 118(12):e2021745118. doi:10.1073/pnas.2021745118.

Parker, David B., and Qolamreza R. Razlighi. 2019. “The Benefit of Slice Timing Correction in Common fMRI Preprocessing Pipelines.” Frontiers in Neuroscience 13. doi:10.3389/fnins.2019.00821.

Pessoa, Luiz, and Jan B. Engelmann. 2010. “Embedding Reward Signals into Perception and Cognition.” Frontiers in Neuroscience 4. doi:10.3389/fnins.2010.00017.

Polo, Pablo, Gabriela Fajardo, Jose Antonio Muñoz-Reyes, Nohelia T. Valenzuela, Montserrat Belinchón, Oriana Figueroa, Ana Fernández-Martínez, Marcel Deglín, and Miguel Pita. 2024. “The Role of Exogenous Testosterone and Social Environment on the Expression of Sociosexuality and Status-Seeking Behaviors in Young Chilean Men.” Hormones and Behavior 161:105522. doi:10.1016/j.yhbeh.2024.105522.

Poser, Benedikt A., Maarten J. Versluis, Johannes M. Hoogduin, and David G. Norris. 2006. “BOLD Contrast Sensitivity Enhancement and Artifact Reduction with Multiecho EPI: Parallel-acquired Inhomogeneity-desensitized fMRI.” Magnetic Resonance in Medicine 55(6):1227–35. doi:10.1002/mrm.20900.

Power, Jonathan D., Bradley L. Schlaggar, and Steven E. Petersen. 2015. “Recent Progress and Outstanding Issues in Motion Correction in Resting State fMRI.” NeuroImage 105:536–51. doi:10.1016/j.neuroimage.2014.10.044.

Radke, Sina, Inge Volman, Pranjal Mehta, Veerle van Son, Dorien Enter, Alan Sanfey, Ivan Toni, Ellen R. A. de Bruijn, and Karin Roelofs. 2015. “Testosterone Biases the Amygdala toward Social Threat Approach.” Science Advances 1(5):e1400074. doi:10.1126/sciadv.1400074.

Rilling, James K., Alan G. Sanfey, Jessica A. Aronson, Leigh E. Nystrom, and Jonathan D. Cohen. 2004. “The Neural Correlates of Theory of Mind within Interpersonal Interactions.” NeuroImage 22(4):1694–1703. doi:10.1016/j.neuroimage.2004.04.015.

Ruff, Christian C., and Ernst Fehr. 2014. “The Neurobiology of Rewards and Values in Social Decision Making.” Nature Reviews Neuroscience 15(8):549–62. doi:10.1038/nrn3776.

Sapolsky, Robert M. 1991. “Testicular Function, Social Rank and Personality among Wild Baboons.” Psychoneuroendocrinology 16(4):281–93. doi:10.1016/0306-4530(91)90015-L.

Schurz, Matthias, Joaquim Radua, Markus Aichhorn, Fabio Richlan, and Josef Perner. 2014. “Fractionating Theory of Mind: A Meta-Analysis of Functional Brain Imaging Studies.” Neuroscience & Biobehavioral Reviews 42:9–34. doi:10.1016/j.neubiorev.2014.01.009.

Singmann, Henrik, and David Kellen. 2019. “An Introduction to Mixed Models for Experimental Psychology.” in New Methods in Cognitive Psychology. Routledge.

Tuiten, Adriaan, Jack Van Honk, Hans Koppeschaar, Coen Bernaards, Jos Thijssen, and Rien Verbaten. 2000. “Time Course of Effects of Testosterone Administration on Sexual Arousal in Women.” Archives of General Psychiatry 57(2):149–53. doi:10.1001/archpsyc.57.2.149.

Van Overwalle, Frank. 2009. “Social Cognition and the Brain: A Meta-Analysis.” Human Brain Mapping 30(3):829–58. doi:10.1002/hbm.20547.

Viswanathan, Shivakumar, Rouhollah O. Abdollahi, Bin A. Wang, Christian Grefkes, Gereon R. Fink, and Silvia Daun. 2020. “A Response-locking Protocol to Boost Sensitivity for FMRI -based Neurochronometry.” Human Brain Mapping 41(12):3420–38. doi:10.1002/hbm.25026.

Yarkoni, Tal, Deanna M. Barch, Jeremy R. Gray, Thomas E. Conturo, and Todd S. Braver. 2009. “BOLD Correlates of Trial-by-Trial Reaction Time Variability in Gray and White Matter: A Multi-Study fMRI Analysis” edited by B. Baune. PLoS ONE 4(1):e4257. doi:10.1371/journal.pone.0004257.

Zink, Caroline F., Yunxia Tong, Qiang Chen, Danielle S. Bassett, Jason L. Stein, and Andreas Meyer-Lindenberg. 2008. “Know Your Place: Neural Processing of Social Hierarchy in Humans.” Neuron 58(2):273–83. doi:10.1016/j.neuron.2008.01.025.

